# Nonnegative matrix factorization integrates single-cell multi-omic datasets with partially overlapping features

**DOI:** 10.1101/2021.04.09.439160

**Authors:** April R. Kriebel, Joshua D. Welch

## Abstract

Single-cell genomic technologies provide an unprecedented opportunity to define molecular cell types in a data-driven fashion, but present unique data integration challenges. Integration analyses often involve datasets with partially overlapping features, including both shared features that occur in all datasets and features exclusive to a single experiment. Previous computational integration approaches require that the input matrices share the same number of either genes or cells, and thus can use only shared features. To address this limitation, we derive a novel nonnegative matrix factorization algorithm for integrating single-cell datasets containing both shared and unshared features. The key advance is incorporating an additional metagene matrix that allows unshared features to inform the factorization. We demonstrate that incorporating unshared features significantly improves integration of single-cell RNA-seq, spatial transcriptomic, SHARE-seq, and cross-species datasets. We have incorporated the UINMF algorithm into the open-source LIGER R package (https://github.com/welch-lab/liger).

## Introduction

Each cell type or state within an organism is distinguished by its gene expression, epigenetic regulation, and spatial location within a tissue. Single-cell sequencing technologies measure each of these features in individual cells, allowing researchers to classify cells in a data-driven manner. Determining what features are common, or different, between each cell type provides researchers insight into the function of the cell. Comparing the profiles of diseased cells with those of healthy cells also reveals disease-related aberrant features. An ideal characterization of a cell type goes beyond analyzing features such as epigenetic and gene expression individually, instead examining their relationships. Jointly examining cellular features holds promise for understanding gene regulatory mechanisms that control cell fates.

Current single-cell sequencing technologies cannot simultaneously measure all relevant aspects of cell state. In particular, emerging techniques such as the Multiome assay from 10X Genomics can measure gene expression and chromatin accessibility from the same cell^1–6^, but do not generally capture methylation or spatial features. Spatial transcriptomics, named Method of the Year 2020 by Nature Methods^7^, encompasses a rapidly growing suite of techniques^8–11^ that interrogate gene expression patterns within intact tissue. However, protocols for spatial measurements of epigenomic state are not widely available.

The different types of features measured by different single-cell technologies create unique computational data integration challenges. Most existing computational approaches for multi-omic data integration are designed for either vertical or horizontal integration scenarios^12^.

Vertical integration approaches are useful for datasets measured across a common set of samples or cells. Well-established methods for multi-omic integration of bulk data such as similarity network fusion and iCluster^13,14^ fall into this category, as well as recent methods for single-cell datasets with multiple modalities per cell such as MOFA+, totalVI, and the Seurat weighted nearest neighbors algorithm^15–17^. Conversely, horizontal integration uses a set of common variables or features to integrate over multiple experiments, typically using shared genes as the basis of integration. Batch effect correction approaches originally designed for bulk sequencing data (e.g., RUV^18,19^ and ComBat^20^) solve a horizontal integration problem. Similarly, “dataset alignment” algorithms developed for single-cell data, such as Seurat, Harmony, and our previous method LIGER^15,21,22^ also rely on shared features and can thus be considered horizontal integration techniques.

In short, LIGER, as well as Seurat and Harmony, is constrained to integrate across features shared between datasets. These methods require that the input matrices all contain a common set of genes or features that are measured in all datasets. Thus, these methods cannot incorporate features unique to one or more datasets, such as intergenic epigenomic information.

Restricting single-cell integration analyses to features shared across all datasets is problematic because it often necessitates discarding pertinent information. For instance, scRNA-seq measures transcriptome-wide gene expression within individual cells, but spatial transcriptomic protocols often measure only a chosen subset of all genes. Yet for many applications, we want to integrate scRNA-seq and spatial transcriptomic datasets, which have neither the same number of features (genes) nor the same number of observations (cells). By integrating the datasets using only shared features, we fail to capitalize on the higher resolution provided by the scRNA-seq modality. When integrating cross-species datasets, the integration is restricted to homologous genes, disregarding all genes without unambigious one-to-one relationships between species. Likewise, when integrating single-cell epigenomic data with single-cell transcriptomic data, horizontal integration approaches do not take into account the important epigenomic features from intergenic regulatory elements. As a final example, existing methods do not provide a way to leverage paired epigenomic information when integrating data types such as SNARE-seq^1,4^ or 10X Multiome with single-cell or spatial transcriptomic datasets. Such integration analyses do not fit neatly into either the horizontal or vertical integration paradigm, requiring the development of new methods.

The critical need to include unshared features in single-cell integration analyses motivated us to extend our previous approach. We developed UINMF, a novel nonnegative matrix factorization algorithm that allows the inclusion of both shared and unshared features. UINMF can integrate data matrices with neither the same number of features (e.g., genes, peaks, or bins) nor the same number of observations (cells). Furthermore, UINMF does not require any information about the correspondence between shared and unshared features, such as links between genes and intergenic peaks. By incorporating unshared features, UINMF fully utilizes the available data when estimating metagenes and matrix factors, significantly improving sensitivity for resolving cellular distinctions.

## Results

### New Integrative Nonnegative Matrix Factorization Algorithm for Partially Overlapping Feature Sets

The key innovation of UINMF is the introduction of an unshared metagene matrix *U* to the iNMF objective function, incorporating features that belong to only one or a subset of the datasets when estimating metagenes and cell factor loadings. Previously, dataset integration using the iNMF algorithm operated only on features common to all datasets. Each dataset (*E*_*i*_) was decomposed into dataset specific metagenes (*V*_*i*_), shared metagenes (*W*), and cell factor loadings (*H*_*i*_), and the optimization problem was solved iteratively. By including an unshared metagene matrix (*U*_*i*_), we provide the capability to include unshared features during each iteration of the optimization algorithm (**Fig. 1a**). The new optimization algorithm does not introduce significant computational complexity compared to our previous iNMF algorithm (see **Methods**).

**Figure 1:**
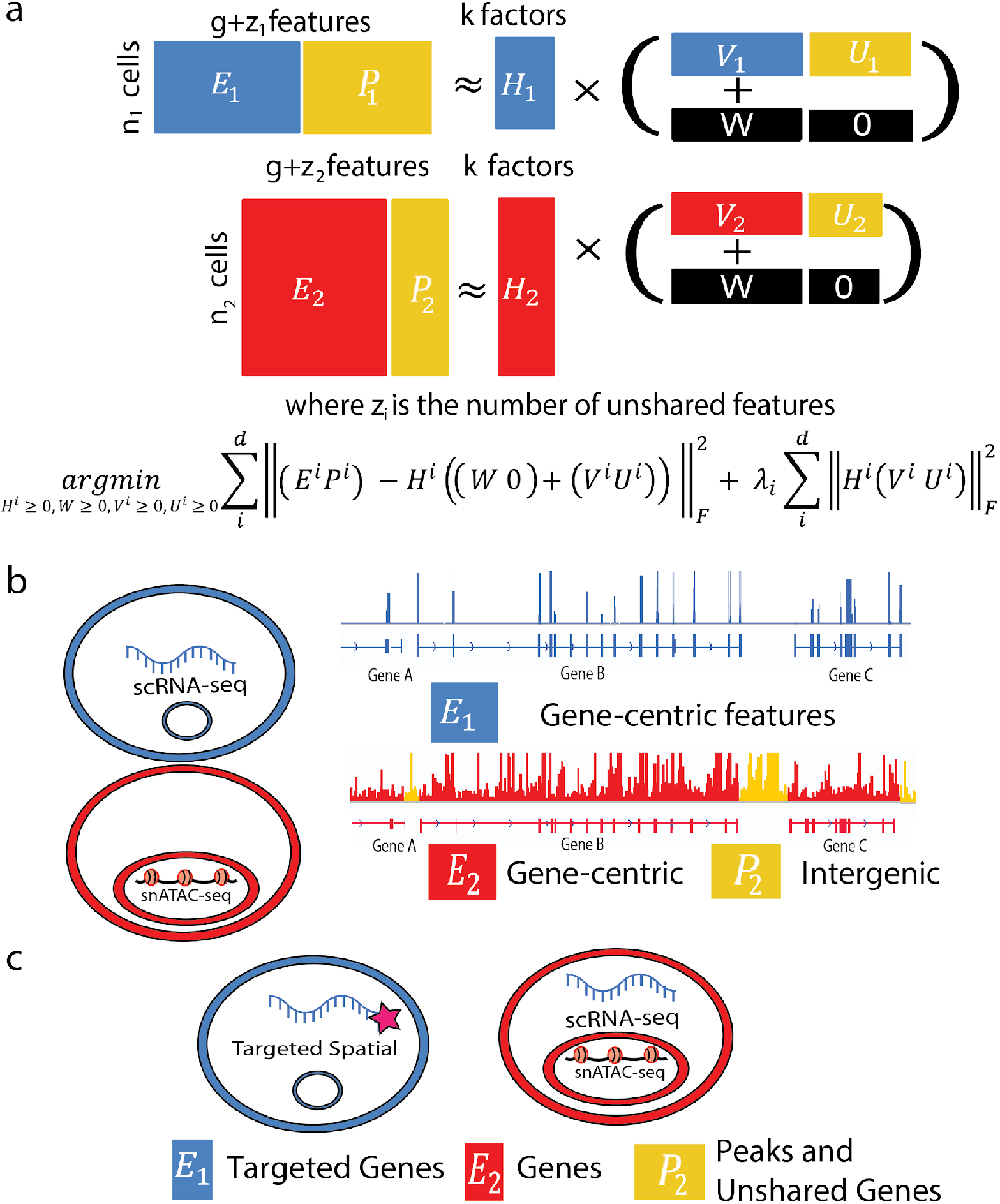
Overview of new nonnegative matrix factorization algorithm for integrating single-cell datasets with partially overlapping features. **(a)** Schematic representation of matrix factorization strategy (top) and optimization problem formulation (bottom). The addition of matrix *U*_*i*_ allows for unshared features to be utilized in joint matrix factorization. Each dataset (*E*_*i*_) is decomposed into shared metagenes (*W*), dataset-specific metagenes (V_*i*_), unique metagenes (*U*_*i*_), and cell factor loadings (*H*_*i*_). The incorporation of the *U* matrix allows features that occur in only one dataset to inform the resulting integration. **(b)** UINMF can integrate data types such as scRNA-seq and snATAC-seq using both gene-centric features and intergenic information. **(c)** UINMF can integrate targeted spatial transcriptomic with simultaneous single-cell RNA and chromatin accessibility measurements using both unshared epigenomic information and unshared genes.

The unshared feature matrix can include extra genes, intergenic features, non-homologous genes, or any other data type that is measured in one of the datasets. Importantly, UINMF makes no assumptions about the relationship between the unshared features and the shared features; for example, no prior knowledge about linkages between intergenic peaks and genes is required. Instead, such covariance among features is learned during the optimization process, as both shared and unshared features contribute to the reconstruction of the original data through the inferred latent factors. These properties allow for the use of the UINMF algorithm across diverse contexts. For example, *U*_*i*_ can incorporate intergenic information when integrating single cell transcriptomic and epigenetic datasets (**Fig. 1b**). When analyzing spatial transcriptomic datasets measuring only a few targeted genes, *U*_*i*_ can be used for genes that are not in the targeted panel but are measured in transcriptome-wide scRNA-seq data. With the advent of multi-omic datasets, the flexible nature of *U*_*i*_ is a particular advantage. We can jointly integrate single-modality data with multimodal data, using all of the features present in the multimodal dataset when performing the integration. In this scenario, the *U*_*i*_ matrix can be used for the unshared multimodal data type, such as chromatin accessibility, when integrating SNARE-seq data with spatial transcriptomic or scRNA data (**Fig. 1c**). Another application of the unshared matrix is the ability to include non-homologous genes into cross-species analysis, leveraging species-specific genes in dataset integration. We demonstrate the functionality of the UINMF algorithm in each of these four possible scenarios, and anticipate that the approach will prove useful for a wide variety of future applications.

### Including Intergenic Peak Information Improves Integration of scRNA and snATAC Datasets

We first investigated how the inclusion of additional features might impact integration of scRNA and snATAC datasets. In our previous work, we summed ATAC reads that fell within a gene to provide gene-centric ATAC profiles, then used these shared features for integration, neglecting intergenic information. In contrast, UINMF uses the unshared feature matrix to incorporate the ATAC reads present between genes—intergenic peaks—when estimating the metagenes (**Fig. 2a)**. Single-nucleus ATAC-seq datasets are extremely sparse, with only 1-10% of peaks detected per cell, compared to the 10-45% of genes captured per cell in scRNA-seq methods^23^. Including the intergenic ATAC data allows more of the detected regions—those not associated with any gene—to be used in each cell, a distinct advantage in such sparse datasets. Additionally, the intergenic chromatin peaks provide information about the chromatin state of important *cis*-regulatory elements, such as promoters and enhancers. We hypothesized that the inclusion of this additional information would better resolve molecular distinctions among cells when integrating single-cell transcriptomic and epigenomic datasets.

**Figure 2.**
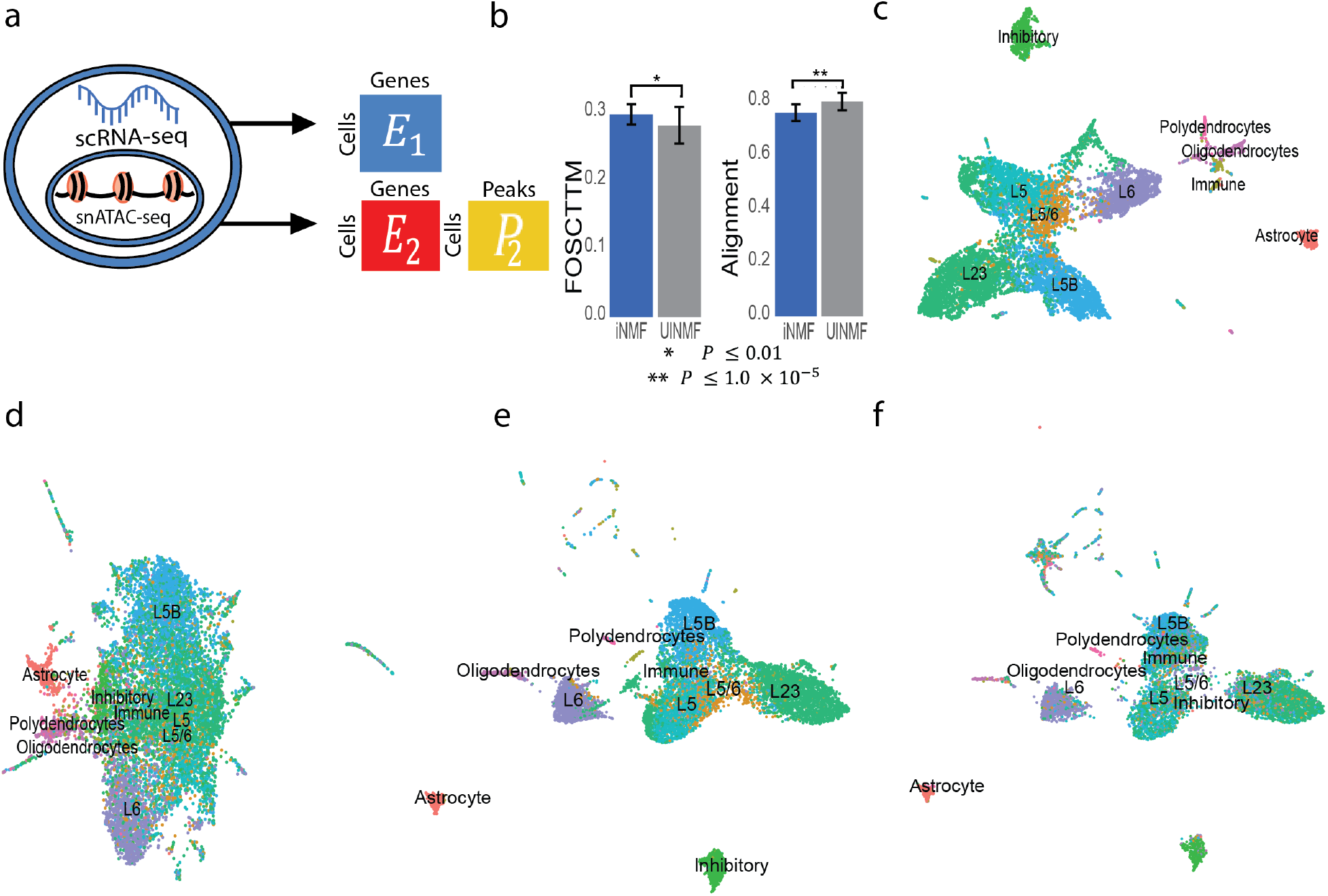
Addition of Intergenic Peak Information Improves Integration of RNA and ATAC Datasets. **(a)** Schematic illustrating how the UINMF algorithm incorporates intergenic peaks when separately integrating the RNA and ATAC measurements from a SNARE-seq dataset. We treat each data type as if it came from an independent source, and perform an integration using regular iNMF and our proposed UINMF method, which incorporates intergenic peaks. **(b)** Average FOSCTTM and alignment scores over thirty initializations of the two algorithms. UINMF achieves significantly better values of both metrics. We factorize and cluster the cells using their RNA transcripts **(c)** and chromatin accessibility measures **(d)** separately. After integration, we use the known cell correspondences to separately plot the gene expression **(e)** and chromatin accessibility datasets **(f)** from SNARE-seq, colored by the same cell type labels.

To quantify how leveraging intergenic features improves dataset integration, we analyzed a SNARE-seq dataset^1^, which provides gene expression and chromatin accessibility information from the same barcoded cell. Because the RNA and ATAC information is measured within the same single cells, the joint profiles provide ground truth cell correspondence information for assessing integration performance. The RNA and ATAC profiles can be preprocessed and integrated as if they come from separate datasets. Subsequently, the success of the integration can be measured by how closely the ATAC and RNA profiles for the same cell are aligned.

We evaluated the quality of SNARE-seq integrations using the Fraction of Samples Closer Than the True Match (FOSCTTM)^2^ metric. The FOSCTTM metric assesses how closely the RNA barcoded cell is placed to its corresponding ATAC barcode in the latent space. Lower FOSCTTM scores are better, indicating that the RNA and ATAC profiles from the same cells have been correctly placed near each other. We also calculated an alignment metric; because the RNA and ATAC datasets come from identical cells, perfect alignment is theoretically achievable and thus the ideal performance is an alignment score of 1.

We assessed the benefit of incorporating intergenic peaks by comparing the UINMF algorithm with our previously published iNMF algorithm^22^, which uses only gene-centric features. Over multiple random initializations, iNMF obtained an average FOSCTTM score of 0.2984 and an average alignment score of 0.761 (**Fig. 2b**).

In contrast, UINMF achieved a significantly lower average FOSCTTM score of 0.2822 (*P* = 0.0064, paired one-sided Student T-test), as well as a significantly higher average alignment score of 0.8005 (*P* = 6.945 × 10^−6^, paired one-sided Student T-test). These findings indicate that incorporating unshared features in the chromatin accessibility data improves the integration of scRNA and scATAC datasets.

We also confirmed that the RNA and ATAC profiles were mapped to similar cell types. To do this, we manually annotated the cell type labels using marker genes from the scRNA data only. Before integration, the clusters separated much more clearly from gene expression data alone than from chromatin accessibility data alone (**Fig. 2 c-d**). After UINMF integration, the cluster labels aligned well across datasets (**Fig. 2 e-f**), indicating that UINMF identified corresponding cell types even though the single-cell correspondence information was not used by the algorithm. In summary, including intergenic chromatin accessibility information in the integration of scRNA and snATAC data significantly improved the integration results.

### Leveraging Additional Genes Improves Integration with Targeted Spatial Transcriptomic Technologies

We expect the UINMF algorithm to be especially effective for integrating targeted spatial transcriptomic datasets, as the number of genes measured in such datasets is often limited. Integrating such datasets with scRNA-seq provides the opportunity to pair transcriptome-wide profiles from dissociated cells with spatial transcriptomic data. This mitigates loss of sensitivity for distinguishing cell types while mapping cell types to their spatial positions within a tissue.

To explore the utility of the UINMF algorithm for integrating targeted spatial transcriptomics and scRNA-seq datasets, we analyzed STARmap data^9^ and scRNA-seq data^24^. We used a STARmap dataset that contains spatial position and transcription level for 28 genes across 31,294 cells within a 3D block of tissue from the mouse frontal cortex. To our knowledge, this dataset is unique in that it is the only dataset of spatially resolved gene expression for multiple genes within a 3D tissue block.

Even though the 28 genes are selected to distinguish among cell types, the STARmap data fails to separate cortical cell types as clearly as scRNA-seq data. We observe improved cluster resolution using iNMF with only shared genes to integrate the STARmap dataset with the scRNA-seq dataset from Saunders (**Fig. 3c**), but some distinct cell types are still mixed together, while others are arbitrarily split. The original cluster labels of the scRNA-seq data are not very well-preserved in the resulting clusters. For example, no distinct boundaries are apparent between the Layer 6, Layer 5, and Layer 5B excitatory neurons. The mural cells are likewise ill-defined. Using UINMF to incorporate 3,525 more genes into the integration allows the metagenes to be estimated from a broader array of genes. Consequently, the addition of these unshared features results in dramatically clearer clusters that much better reflect the ground truth labels (**Fig. 3d**). The distinction between the excitatory neurons subtypes becomes clear, and a defined population of mural cells also becomes distinguishable. Thus, using the unshared features, it is possible to identify cell types that would not be otherwise distinguishable.

**Figure 3.**
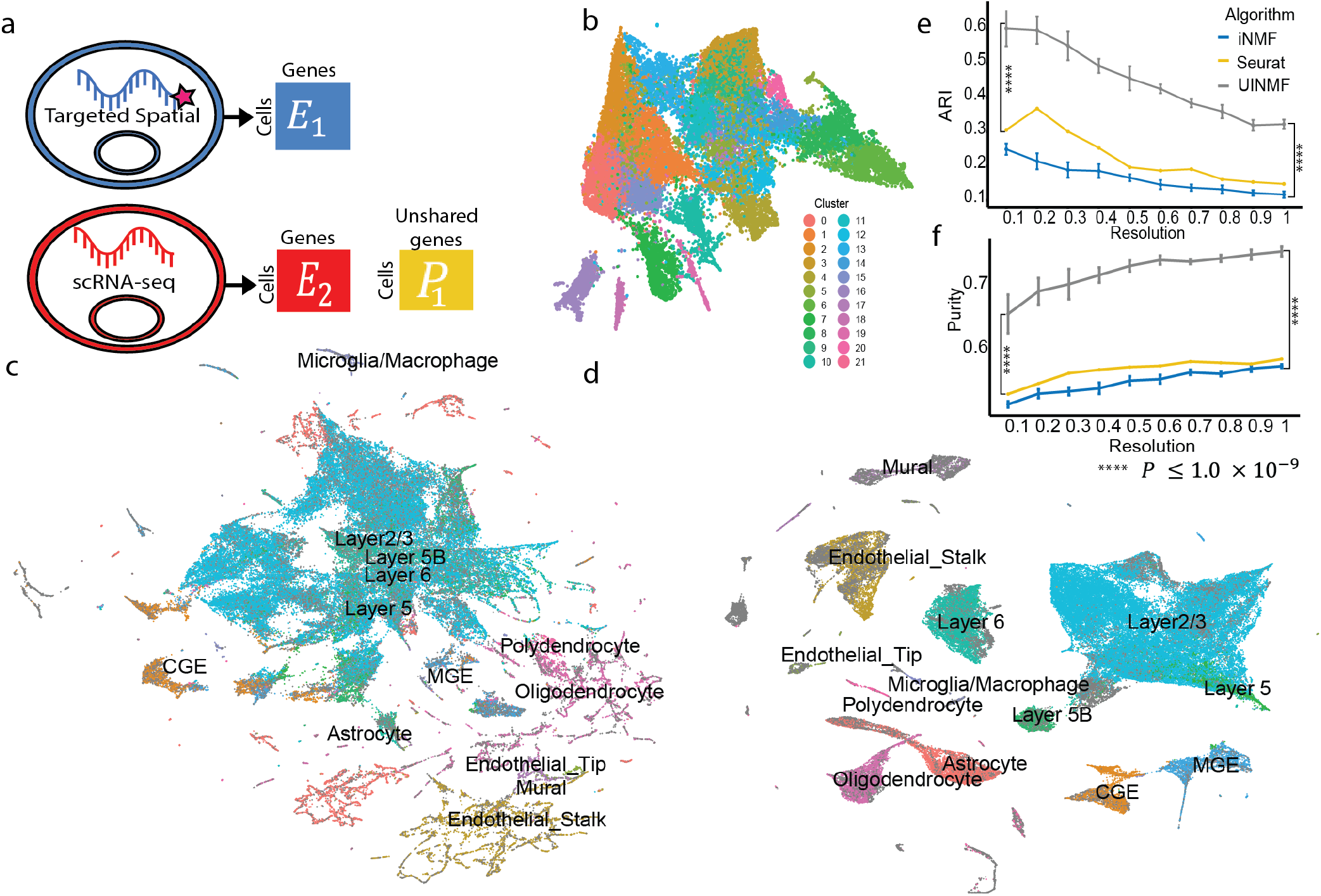
Incorporating additional genes improves integration with STARmap data. **(a)** Schematic of UINMF integration of the spatial transcriptomic data with the scRNA-seq data, in which *U* incorporates unshared genes that are captured in scRNA-seq but not targeted STARmap data. **(b)** UMAP of STARmap data alone. **(c)** UMAP of STARmap and scRNA integration performed with iNMF using only shared genes. **(d)** UMAP of UINMF integration, which incorporates both shared and unshared genes. **(e)**-**(f)** Comparison of adjusted rand index (e) and purity (f) metrics from iNMF, Seurat, and UINMF for a range of clustering resolutions.

To quantify the advantage of the UINMF method, we calculated Adjusted Rand Index (ARI) and purity metrics for multiple initializations of the UINMF and iNMF algorithms across a range of different Louvain resolution parameters (**Fig. 3e,f**). The UINMF algorithm achieves significantly (*P* = 3.895 × 10^−10^, paired one-sided Wilcoxon test) higher ARI and cluster purity compared to the iNMF algorithm across the range of resolution parameters. UINMF also significantly outperforms Seurat in both ARI and purity metrics (*P* = 3.895 × 10^−10^, paired one-sided Wilcoxon test). In short, the addition of the unshared genes significantly improves the integration of STARmap and single-cell RNA-seq data.

To verify that corresponding cell types were clustered together across technologies, we examined the expression of several key marker genes for each labeled cell type by dataset (**Fig. 4a**). Generally, marker genes highly elevated in the cell types of the STARmap dataset are also highly elevated in the corresponding scRNA-seq cell type, indicating that the metagene definitions reflect biological distinctions significant for both datasets. The cell clusterings are not reflective of technical artifacts specific to an individual dataset, nor are they formed overwhelmingly by a single dataset. Rather, the clusters reflect divisions significant to both datasets jointly. Additionally, as we previously noted^22^, there is evidence that the STARmap gene capture is somewhat non-specific compared to scRNA-seq for some genes, such as *Sulf2* and *Mgp*.

**Figure 4.**
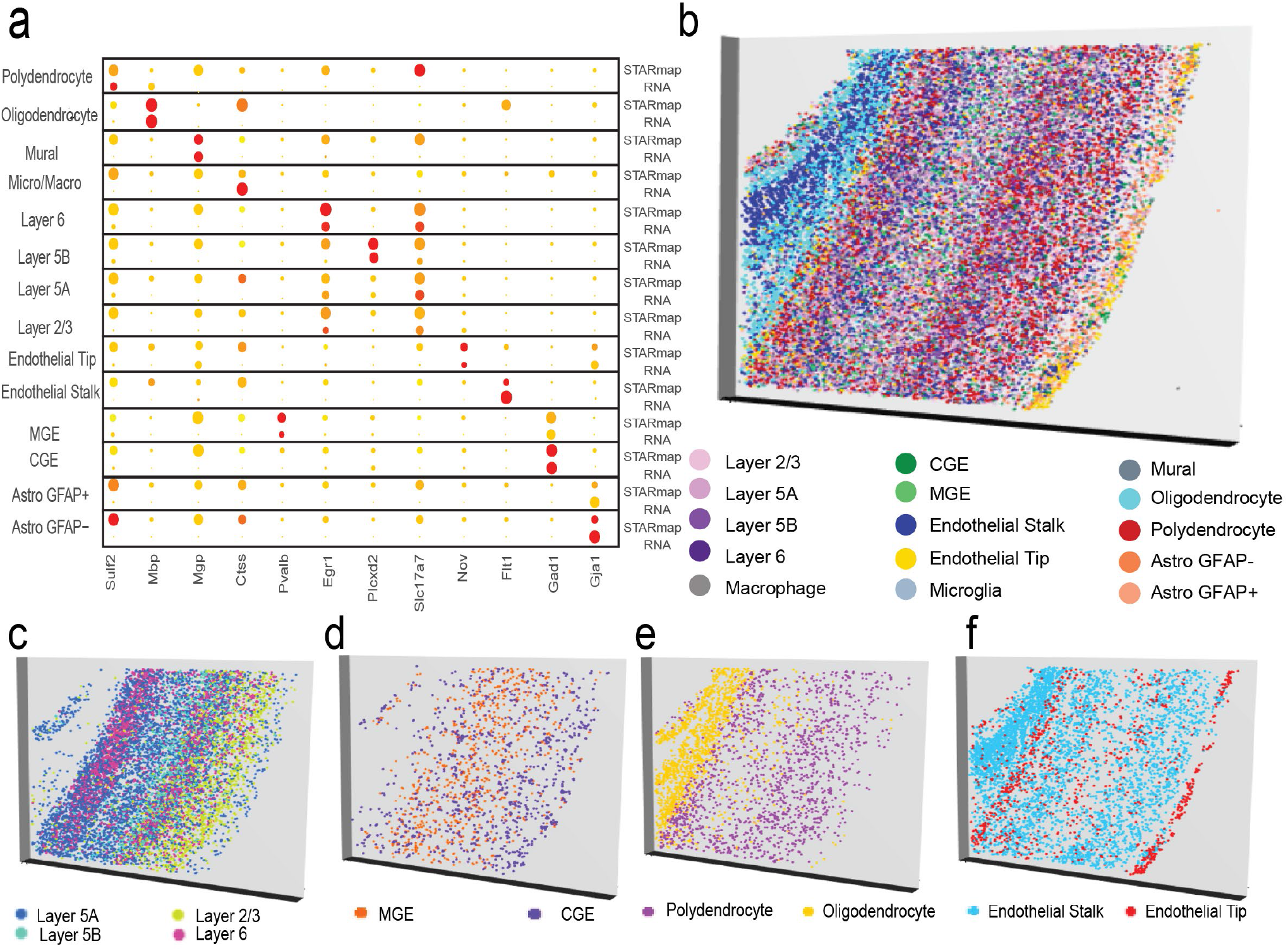
Incorporating additional genes locates fine cellular subtypes within 3D spatial volume. **(a)** Dot plot showing marker gene expression in STARmap and scRNA datasets for each joint cluster. The datasets have similar marker expression, indicating that they are well aligned. Plots of three-dimensional spatial locations for different classes of cells colored by cell type: **(b)** all cell types; **(c)** excitatory neurons; **(d)** inhibitory neurons; **(e)** polydendrocytes and oligodendrocytes; and **(f)** endothelial cells.

A key advantage of refining cell types in the STARmap dataset is the ability to use these cell labels within a 3D tissue sample, providing greater insight about how transcriptomic profiles are arranged in vivo. Consequently, we used the newly derived cell type labels within the context of 3D space, allowing us to assess their validity on the basis of concordance with known tissue architecture (**Fig. 4b**). This analysis confirms that our cell type annotations accord well with the known structure of the cortex, such as clear laminar arrangement of excitatory neurons (**Fig. 4c**). It has also been previously established that MGE interneurons originate from the more rostral region of the brain, and the CGE neuron center lies caudal to the MGE center^25^. Likewise, the MGE and CGE determined cell types establish a gradient such that the CGE interneurons increase as in more cranial regions of the cortex slice (**Fig. 4d**). The region of white matter that lies beneath the cortex, composed primarily of oligodendrocyte cells, is also identifiable (**Fig. 4e**). Playing a significant role in supporting the brain’s vascular systems (7), endothelial cells compose portions of the blood brain barrier, and the UINMF results indicate that endothelial tip cells are located near the outer surface of the brain (**Fig. 4f**).

We further demonstrate the advantages of UINMF for spatial transcriptomic datasets by integrating the osmFISH dataset (5,185 cells and 33 genes)^11^ from the mouse somatosensory cortex with the DROPviz scRNA-seq dataset from frontal cortex (71,639 cells). Using iNMF to integrate the osmFISH and DROPviz datasets utilizes the 33 genes measured in the osmFISH protocol as the shared features (**Fig. 5a**). The iNMF integration successfully aligns the two data types, but, similar to the STARmap analysis, leads to both mixing and oversplitting of cell types. The intermixing of cell types, especially the excitatory neurons, fails to resolve a distinct cluster for the Layer 5B Excitatory neurons. Using UINMF to incorporate an additional 2,000 variable genes from the scRNA-seq dataset allows the metagenes to be estimated from many more features, resulting in much more clearly resolved clusters (**Fig. 5c**). A distinct cluster of Layer 5B Excitatory neurons can now be distinguished. Including additional features into the integration not only categorizes broad cell types more effectively, but also identifies more minute subclasses of cells.

**Figure 5.**
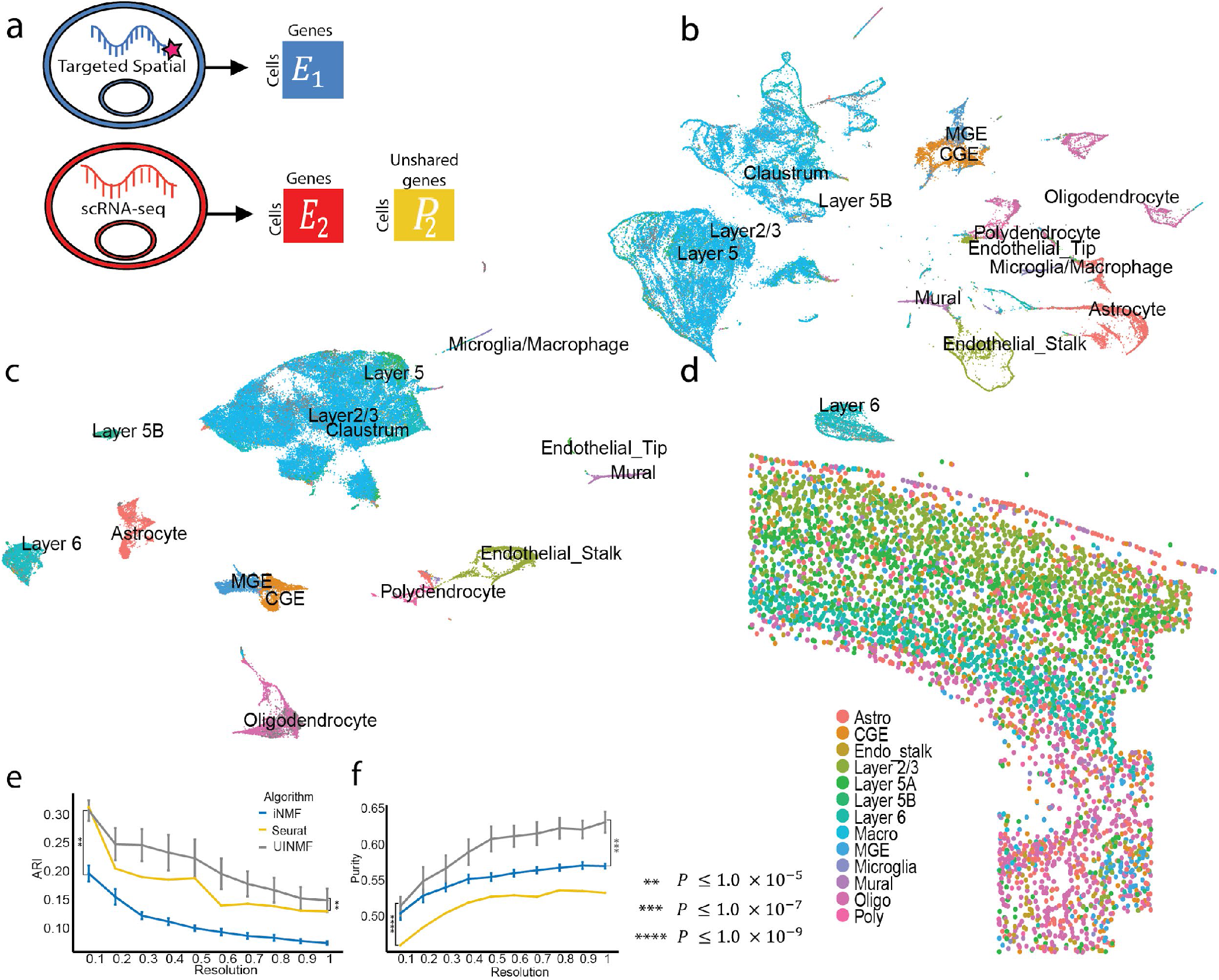
Incorporating additional genes improves integration with osmFISH data. **(a)** Schematic of data matrices from osmFISH and scRNA-seq. The osmFISH dataset measures only 33 genes, while the scRNA-seq dataset has many unshared genes that are incorporated during UINMF integration. **(b)** UMAP plot of osmFISH and scRNA integration with iNMF using only shared genes. **(c)** UMAP using UINMF to incorporate an additional 2,000 genes. **(d)** The spatial arrangement of cell types matches the known tissue structure of the cortex **(e)-(f)** ARI (e) and purity (f) metrics for iNMF, UINMF, and Seurat.

As with the STARmap data, we then plotted the UINMF labels within their corresponding spatial context (**Fig. 5d**). The excitatory cells again show clear laminar arrangement, with layer 6 excitatory neurons forming the innermost layer of the cortex, and layers 5 and layer 2/3 neurons above. Interestingly, layer one contains a number of cells identified as astrocytes, a finding that has previously been observed experimentally and which has been proposed as evidence for the interaction of glial cells in neuronal signaling^26^. The white matter region, located inferior to layer 6 and known to be composed of oligodendrocytes and polydendrocytes^27,28^, is likewise observable. Lastly, the presence of the caudoputamen and internal capsula region can be identified by the small grouping of inhibitory neurons lateral to the white matter^11^.

To quantify the advantage of using UINMF, we measured the ARI and purity scores for both UINMF and iNMF over multiple initializations and multiple clustering resolutions (**Fig. 5 e**,**f**). UINMF performed significantly in terms of both ARI (*P* < 2.2 × 10^−16^, paired one-sided Wilcoxon test) and purity (*P* = 7.078 × 10^−8^paired one-sided Wilcoxon test) metrics across the whole range of clustering resolutions. A similar improvement was observed when comparing the UINMF performance to Seurat. UINMF performed significantly better in both ARI (*P* = 3.946 × 10^−6^)and purity (*P* < 2.2 × 10^−16^) metrics.

### UINMF Improves Integration of Multimodal and Spatial Transcriptomic Datasets

Single-cell multimodal technologies measure epigenomic and transcriptomic profiles from the same cell, providing an exciting opportunity to define cell types from multiple molecular modalities. However, many applications require integrating such multimodal measurements with single-modality datasets. In such applications, the ability of UINMF to incorporate unshared features allows us to capitalize on the multimodal information, rather than using only shared features.

To demonstrate the advantages of such an approach we used UINMF to integrate STARmap and SNARE-seq data. The STARmap dataset provides gene expression data for 2,522 cells across 1,020 genes while preserving the 2D spatial coordinates for each cell. The SNARE-seq dataset (10,309 cells) provides simultaneous chromatin accessibility and gene expression levels from the same barcoded cells.

We first integrated the STARmap data and SNARE-seq gene expression measurements only by performing iNMF on the 944 genes shared between the datasets, omitting the unshared genes and completely neglecting the chromatin information. We annotated the cells by using the original annotations from both datasets to jointly define the resulting clusters (**Fig. 6a**). When integrating the datasets with UINMF, we were able to add an additional 2,688 highly variable genes present in the SNARE-seq dataset. Because the SNARE-seq data is multi-omic, we also incorporated the available chromatin accessibility information by including the top 1,431 variable chromatin accessibility features within the U matrix (**Fig. 6b**). Thus, the UINMF integration incorporated a total of 4,119 features not measured in the STARmap dataset (**Fig. 6c**).

**Figure 6.**
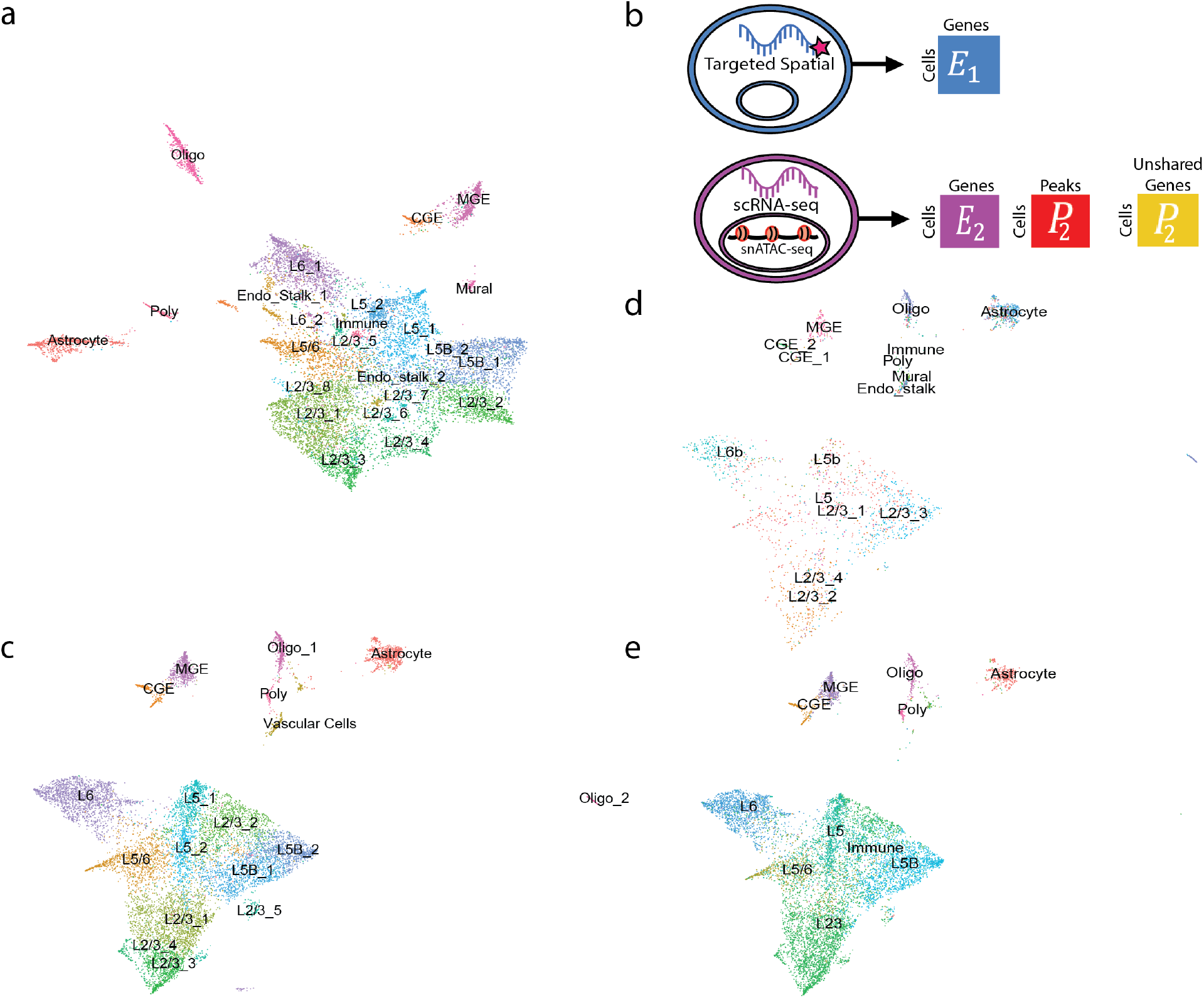
Incorporating unshared chromatin and gene features to integrate spatial transcriptomic and multimodal data. **(a)** UMAP for iNMF integration of STARmap spatial transcriptomic data and SNARE-seq RNA data only. **(b)** Schematic of how unshared gene and chromatin accessibility data is incorporated into the integration analysis of STARmap and SNARE-seq using UINMF. **(c)** The integration is improved significantly by the inclusion of ATAC gene-centric features in the U matrix. **(d)-(e)** The original cell type labels of STARmap cells (d) and SNARE-seq cells (e) show clear correspondence after UINMF integration.

Next, we confirmed that the integrity of each individual dataset had been maintained by examining the STARmap and SNARE-seq cells individually by their original labels (**Fig. 6d,e**). There is clear alignment between the original cell labels of each dataset, indicating that the integration defined the metagenes relevant to specific cell populations for both datasets. This suggests that the unshared features can be included into the integration without unduly dominating the metagene calculations.

To quantify the derived benefit of including the additional features into the analysis, we then calculated the purity and ARI scores for ten initializations (**Fig. 7a,b**). UINMF significantly outperformed the iNMF algorithm on both ARI (*P =* 5.934 × 10^−15^, paired one-sided Wilcoxon test) and Purity (*P* = 1.046 × 10^−11^, paired one-sided Wilcoxon test) and purity. The iNMF algorithm has an ARI competitive with that of UINMF at only a single louvain resolution (0.7). At this resolution, UINMF still has a superior purity score, substantiating the benefit of including additional features using the UINMF algorithm. UINMF also has a significantly better ARI score than Seurat across resolutions (*P* = 0.001562). Additionally, the UINMF also has superior purity over Seurat at all tested resolutions (*P* = 3.016 × 10^−16^, paired one-sided Wilcoxon test). Taken together, these results show that the use of UINMF to incorporate added features to the data integration significantly improves integration of single-cell multimodal data and spatial transcriptomic data.

**Figure 7.**
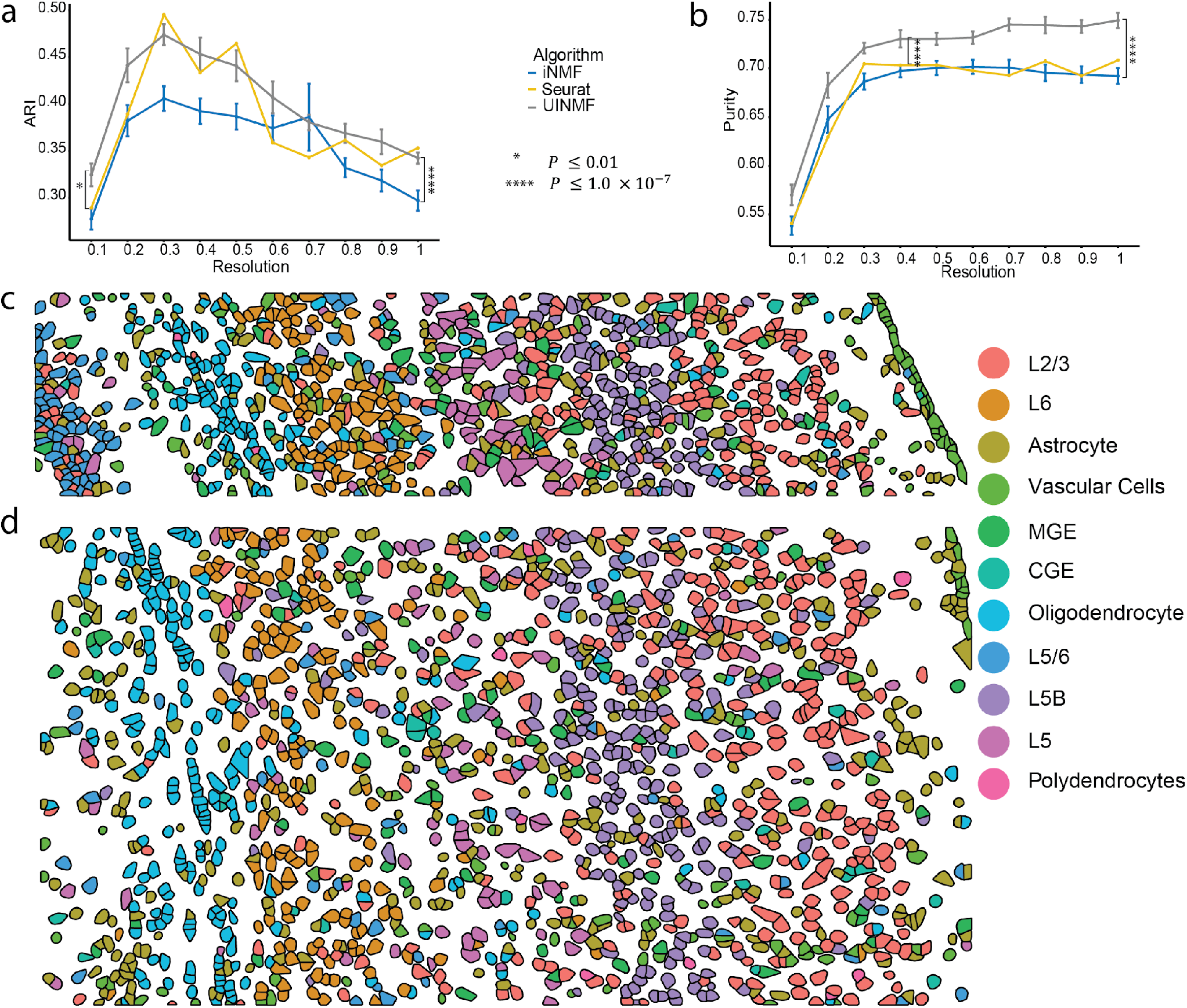
Incorporating unshared chromatin and gene features improves integration of spatial transcriptomic and multimodal data. We compared UINMF results with those from iNMF and Seurat using purity **(a)** and ARI **(b)** metrics. We also confirmed that the spatial arrangement of predicted cell types in both STARmap replicate one **(c)** and replicate two **(d)** matches the known organization of the cortex.

Because a key motivation for integrating the multimodal and spatial transcriptomic data was bringing enhanced resolution within the context of spatial coordinates, we next plotted the results of the UINMF integration in space. Applying the cell type labels from UINMF to STARmap replicate one (973 cells, **Fig. 7c**) and replicate two (1,549 cells, **Fig. 7d**), we found that the UINMF results accord well with the cortical structure. The excitatory neurons are arranged in layers, with L6, L5, and L2/3 clearly visible. Likewise, we also can identify the oligodendrocyte-rich white-matter below the cortex. Additionally, the vascular cell population, which contains endothelial and mural cells comprise the outermost group of cortical cells.

### The Inclusion of Non-Homologous Genes Improves the Integration of Cross-Species Data

Previous integrations of cross-species datasets have been limited to genes that are homologous between species, as non-homologous genes, by definition, are not shared between datasets. Yet, non-homologous genes can be key marker genes within a species, providing crucial information for distinguishing cell populations. Using UINMF, we were able explore the potential benefits of including non-homologous genes in cross-species integration. For this cross-species analysis, we integrated scRNA data (4,187 cells) from the pallium of the bearded dragon lizard (*Pogona vitticeps)*^29^ with scRNA data from the mouse frontal cortex (71,639 cells)^24^. We first selected 1,979 variable genes that were annotated as one-to-one homologs between the two species. Then we selected 166 non-homologous variable genes from the lizard dataset, and an additional 1,110 non-homologous variable genes from the mouse dataset. We integrated the datasets using UINMF, with the one-to-one homologs as shared features, and the non-homologous genes as unshared features (**Fig. 8a**). UINMF successfully aligned the two datasets, as illustrated by the overlapping distributions of the two datasets within the UMAP space (**Fig. 8b**). To confirm correspondence between the cell types of the two species, we plotted only the mouse cells (**Fig. 8c**) and only the lizard cells (**Fig. 8d**), colored by their originally published labels. Strong correspondence between cell types, including excitatory neurons, inhibitory neurons, and non-neuronal cells, can be observed.

**Figure 8.**
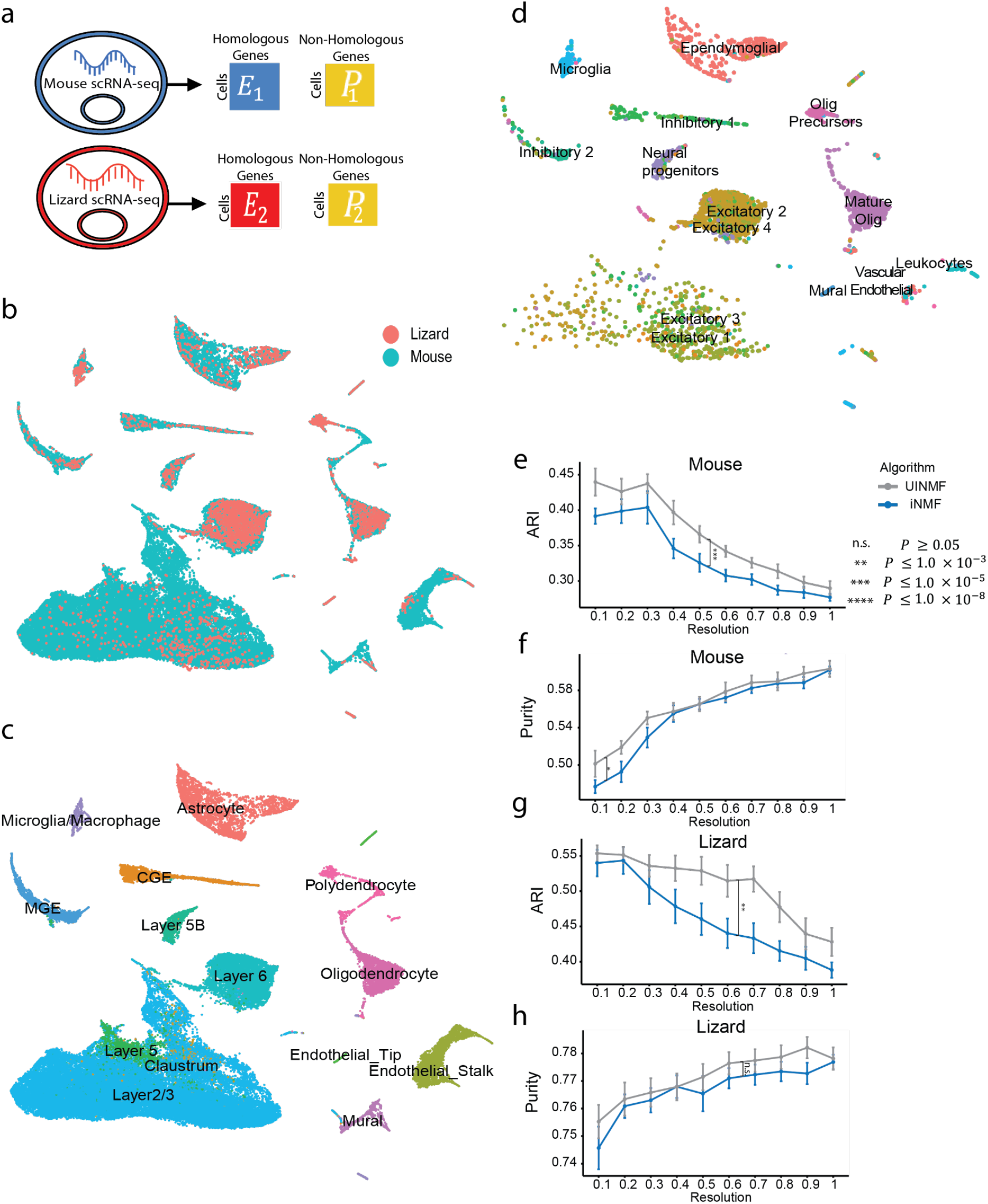
The inclusion of non-homologous genes improves the integration of cross-species data. We demonstrate the alignment between the two datasets **(a)**, after using UINMF to include both homologous and non-homologous genes when integrating the datasets **(b)**. We also confirmed cell type correspondence by examining only the mouse cells **(c)** and only the lizard cells **(d)**, both labeled with their published cell labels. To show the advantage of including the non-homologous genes, we show the difference in ARI **(e)** and purity **(f)** scores using the originally published mouse. We also confirm a similar trend in the ARI **(g)** and purity **(h)** scores using the original lizard labels.

To examine the additional benefit of including non-homologous genes when performing cross-species integration, we performed the integration using iNMF to establish a baseline. The baseline iNMF integration was limited to the 1,979 homologous genes, and resulted in a lower quality integration (**Supplementary Figure 5**). The mural cell populations had decreased alignment between the two species, and many of the astrocytes were misaligned to the excitatory neuron clusters. Furthermore, the lizard’s excitatory neuron subtypes were less distinctly separated.

In order to quantify the advantage of UINMF over iNMF in cross-species integrations, we compared the purity and ARI scores of the two algorithms across ten initializations. Including the non-homologous genes using UINMF resulted in a significant increase in both the ARI *(P =* 3.626 × 10^−9^, paired one-sided Wilcoxon test) and the purity (*P =* 6.258 × 10^−4^, paired one-sided Wilcoxon test) of the mouse dataset (**Fig. 8e,f**). We also noted a significant increase in the ARI (*P =* 1.145 × 10^−6^, paired one-sided Wilcoxon test) of the lizard data set (**Fig. 8g**). Although UINMF does not show a statistically significant increase in the purity (*P =* 0.07157, paired with one-sided Wilcoxon test) of the lizard dataset, UINMF is able to achieve a higher maximum purity scores at most resolutions (**Fig. 8h**). This more modest effect observed for the lizard dataset may be because it had many fewer non-homologous variable genes (166) than the mouse dataset (1,110), likely due to a less complete set of gene annotations for the lizard genome.

## Discussion

We have extended our previous integrative nonnegative matrix factorization algorithm by adding an unshared feature matrix. This addition accommodates features that are not present in all datasets and increases the amount of information that is used to define the metagenes. We showed that inclusion of unshared features provides clear advantage across four different types of integration analyses. First, UINMF can be used to incorporate intergenic information when integrating transcriptomic and epigenomic datasets. Second, UINMF can incorporate genes not measured in targeted spatial transcriptomic datasets, allowing better resolution of fine cellular subtypes within a spatial coordinate frame. Third, UINMF can utilize all of the information present in single-cell multimodal when integrating with single-modality datasets. Additionally, UINMF can accommodate non-homologous genes in cross-species integration analyses.

With the rapid development of multimodal and spatial transcriptomic technologies, we anticipate that the UINMF algorithm will prove useful for a wide variety of analyses. As the additional *U* matrix can incorporate any type of cellular features, with no assumptions about their relationship to gene-centric features, the algorithm is inherently flexible to accommodate a variety of data types and modalities. Future applications could examine the incorporation of data types such as Hi-C^30^ measurements, as well as the potential to use UINMF on a diverse collection of *in situ* hybridization and immunohistochemistry datasets with limited numbers of genes. We expect that, as additional experimental methods for single-cell measurement are developed, our approach will prove increasingly useful for a broad variety of single-cell integration tasks.

## Methods

We extend our previously published ANLS algorithm for solving the iNMF problem^22^ so that we can now incorporate unshared features when integrating across datasets. The unshared feature matrix can in principle accommodate any type of unshared feature, whether gene-centric or otherwise. In this paper, we incorporated intergenic peaks from snATAC-seq data and additional genes not measured in all datasets, although many other applications are possible.

For each data set *E*^1^, *E*^2^,….*E*^*n*^, we normalize the data, and select *m* variable genes (shared across all datasets), and *z*_*i*_ variable features, such that after scaling 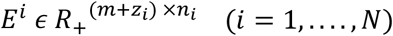. For a given *K* and λ_*i*_, the optimization problem can be defined as

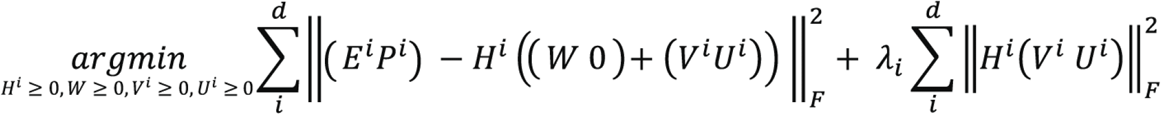

We then solve the UINMF factorization problem with a realization of the coordinate block descent (BCD) approach^31^. The BCD approach divides the parameters into blocks, and then finds the optimal parameters by updating each block while holding the others fixed. Because each block-wise optimization sub-problem is convex, iterating these updates is guaranteed to converge to a local minimum^31^. To solve the UINMF optimization problem, we use matrix blocks, one block for each of 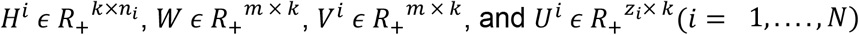. Each sub-problem is a nonnegative least squares optimization, which we solve numerically using an efficient C++ implementation of the block principal pivoting algorithm^32^. To update *H*^*i*^, we solve

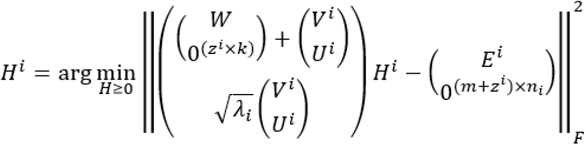

while holding the other parameters fixed. Similarly, to update the other parameters, we solve the following subproblems:

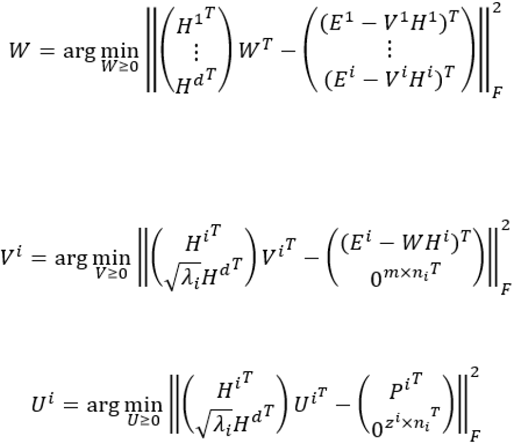

We iterate these updates until convergence, which we determine by calculating the decrease in the objective function at the conclusion of each iteration. We consider the algorithm to have converged when the decrease in the objective value function between the previous and current iteration, weighted by dividing by their mean, is less than the epsilon parameter set by the user. For all UINMF analysis in this paper, we set the convergence threshold to ε = 1.0 × 10^−10^and set the maximum number of iterations to 30 for each analysis.

Note that iNMF uses a constant penalty term λ for all datasets. When implementing UINMF, however, we introduced a separate λ_*i*_ parameter for each dataset, such that the penalty applied to *E* is weighted by λ_1_, *E*_2_ has a regularization weight of λ_2_, etc. The inclusion of λ_*i*_ allows for the tuning of dataset penalization at the user’s discretion. However, we simply used λ_*i*_ = 5 (the default value in our iNMF and UINMF implementations) for all analyses except the STARmap and DROPviz integration. The STARmap and DROPviz integration does achieve the best results with different λ values for each dataset.

### Increase in Computational Complexity

The difference in computational complexity between UINMF and iNMF increases with the number of unshared features used in UINMF, but the difference between the runtime of the two algorithms is not prohibitive in practice (**Supplemental Figure 4**). To assess the theoretical difference in computational complexity between the algorithms, assume the same total number of features is present in datasets input to each algorithm. Let iNMF operate on a dataset that has *g* shared features, and let *g* = *m* + *z*, where *m* is the number of shared features and *z* is the number of unshared features of the UINMF dataset. Let *K* be the user-defined number of metagenes. For each iteration, UINMF solves for *U* ^*z*×*K*^ and *V*_*i*_^*m* × *K*^ separately, but iNMF performs the same number of calculations to solve for *V*_*i*_^*g* × *K*^, since *g* = *m* + *z*. When solving for the shared metagene matrix, *W*, iNMF solves the optimization problem for a *g* × *K* matrix, whereas UINMF must only solve a *m* × *K* matrix. Because the shared metagene matrix has less features in UINMF (*m* < *g*), each iteration of the algorithm actually constitutes less computational complexity than iNMF given the same total number of features.

### Evaluation Metrics

The alignment score, based on Butler et. al (9), is a measure that captures how well two samples align uniformly within a latent space. A score closer to zero indicates a poor alignment, or mixing of the two samples, whereas a score closer to one is indicative of uniformly mixed datasets.

Fraction of Samples Closer Than the True Match (FOSCTTM) scores measures how closely two measurements of the same cell are placed within the latent space^2^. We calculate the FOSCTTM score by finding the distance between the scRNA-seq cell and the scATAC-seq label for each barcode. We then divide by the total number of barcoded cells to derive the average FOSCTTM score.

Cluster purity is calculated based on a reference clustering. Each cluster is assigned a type based on the predominant label for that cluster. The cells that correspond correctly to this label are counted. We calculate purity by summing the correct number of labels across all clusters, and dividing by the total number of labeled cells present. Consequently, a score closer to 0 indicates that the cells are not being accurately grouped into clusters by cell types, and a score of 1 indicates perfect grouping by cell type. Adjusted Rand Index (ARI) is another measure of similarity between two clusterings. The ARI score can range from 0 (no match) to 1 (perfect match).

### Integration of RNA and ATAC Profiles from SNARE-seq

For a baseline, we integrated the scRNA and scATAC datasets using iNMF, using the top 2,589 variable genes and their associated scATAC peaks. The optimization was performed with *K* = 30 and λ = 5. We performed quantile normalization with parameter *knn_k* = 100. We then calculated the average FOSCTTM and alignment scores across thirty random initializations. To assess the additive properties of including the intergenic peaks into the integration, we used the *U*-matrix to hold the top 2,000 variable peaks. We performed UINMF with *K* = 30, λ = 5, and *knn_k* = 20. We calculated FOSCTTM and alignment scores over thirty random initializations. We performed one-sided paired Student T-tests on the resulting FOSCTTM and Alignment scores. To cluster just the RNA and ATAC cells, we similarly used *K* = 30, λ = 5, *knn_k* = 100, and a Louvain resolution of 1. We annotated the cell clusters using marker genes and used the same labels for each UMAP presented in Figure 3.

### Integration of scRNA-seq and STARmap

We integrated the STARmap spatial transcriptomic dataset (32,845 cells) and a scRNA-seq dataset (71,639 cells) using iNMF. For this iNMF analysis, we were limited to the 28 genes measured in the STARmap dataset. Both iNMF and CCA/PCA (used by Seurat) are limited to no more components than the number of genes, while UINMF can estimate more dimensions because it also incorporates unshared genes. We thus used *K* = 27 dimensions for both iNMF and Seurat. For iNMF we also used λ = 5, and a quantile normalization with K-nearest neighbors of 20. Using UINMF, we included an additional 2,775 of the most variable genes. The parameters for the UINMF integration were *K* = 40and λ = 10 for the STARmap data and λ = 1 for the scRNA-seq data; *knn_k*=20 for the quantile normalization; and Louvain resolution of 1. For both iNMF and UINMF, we performed 5 initializations with the same random seed and picked the best one. We calculated the cluster purity for both algorithms using the scRNA-seq labels, and use the highest number of cells present to annotate the clusters by cell type. These annotations were then applied within the context of 3D space using the originally provided STARmap coordinates. For five different random seeds, we measured the difference between the cluster purity (*P* = 3.895 × 10^−10^) and ARI (*P* = 3.895 × 10^−10^) of iNMF and UINMF using a paired, one-sided Wilcoxon test.

For Seurat, we similarly used the 28 shared genes and a PCA dimension of 27. We then calculated the Purity and ARI scores at each Louvain resolution from 0.1 to 1, in increments of 0.1. We performed a paired, one-sided Wilcoxon test to assess the difference between UINMF and Seurat performance quantified by purity (*P* = 3.895 × 10^−10^) and ARI (*P* = 3.895 × 10^−10^).

### Integration of scRNA-seq and osmFISH

Using the osmFISH dataset (33 genes, 6,471 cells) we excluded hippocampal cells and cells from the top left of the tissue slice originally labeled “excluded” in the original osmFISH publication, as well as cells with zero detected genes. A total of 5,185 cells passed these filtering criteria. We ran the iNMF algorithm using the 33 shared genes, λ = 5, and *K* = 32, and took the best optimization of 5 random initializations for each of 10 seeds. Note that, as with STARmap data, iNMF and Seurat are limited to no more components than the number of genes, while UNIMF can estimate more factors due to the use of unshared genes. Using UINMF, we integrated the osmFISH spatial transcriptomic data with the scRNA-seq (71,639 cells) using the 33 shared genes as well as the 2,000 most variable unshared genes. We used 10 different random seeds, with λ = 5 and *K* = 40, and took the best optimization of 5 random initializations for each seed. We calculated the Purity and ARI for each algorithm at Louvain resolutions 0.1 through 1.0, in 0.1 increments. We used the published DROPviz labels as our reference clustering, and assessed the significance of our findings using a paired, one-sided Wilcoxon test, resulting in a significant difference being observed in both purity (*P* = 7.078 × 10^−8^) and ARI score(*P* ≤ 2.2 × 10^−16^). We used *knn_k*=150 for the *quantile_norm* function, *n_neighbors*=150 for the *runUMAP* function, and a resolution of 0.5 for Louvain clustering.

For Seurat integration, we also used the 33 shared genes and a PCA dimension of 32. Using louvain resolutions from 0.1 to 1.0 in 0.1 increments, we assessed the performance of Seurat using purity and ARI scores. Comparing Seurat and UINMF using a paired, one-sided Wilcoxon test showed a significant difference in purity (*P* <= 2.2 × 10^−16^) and in ARI (*P* = 3.946 × 10^−6^).

### Cross-Species Integration

The Lizard Pallium dataset^29^ originally had 4,202 cells, but we limited the cells used to the 4,187 cells deemed high quality in the original publication. We integrated this dataset with the scRNA-seq dataset from the mouse brain^24^ (71,639 cells). We used the original publications 1-to-1 homolog labels to select homologs common to both the mouse and lizard datasets. Using these as our shared features, we normalized and scaled the data. We then selected 1,979 shared genes, using a variance threshold of 0.3. For UINMF, we used the same variance threshold to select 166 non-homologous genes from the lizard, and 1,110 non-homologous genes from the mouse. We optimized UINMF and iNMF with *K* = 30, λ = 5,and took the best of 5 random initializations for 10 random seeds. We performed quantile normalization for each optimized object using the mouse dataset as a reference. To ensure that any differences in purity and ARI score were not driven by a single species, we calculated the ARI and Purity scores using both the lizard and the mouse cell labels separately as ground truth. We performed these calculations for each of the ten seeds at louvain resolutions from 0.1 to 1.0 in increments of 0.1. To examine the difference between UINMF and iNMF performance, we performed a paired, one-sided Wilcoxon test between the ARI (*P =* 3.626 × 10^−9^, *P =* 1.145 × 10−6) and Purity (*P* = 6.258 × 10^−4^, *P =* 0.07157) values, for the mouse and lizard labels, respectively. When generating the UMAPs shown, we used the default louvain resolution of 0.25, and the default nearest neighbors of 10.

### Integration of SNARE-seq and STARmap

To integrate the multi-omic SNARE-seq dataset (10,309 cells) with the spatial transcriptomic STARmap dataset (2,522 cells), we used the number of shared genes (944 genes), as well as 4,119 unshared features. To generate the unshared features, we selected genes with a variance threshold higher than that of 0.1, and then removed genes shared between datasets, for a total number of 2,688 unshared genes. We selected the chromatin accessibility peaks with a variance greater than 0.01 (1,431 peaks). For iNMF benchmarking, we used only the 944 shared genes. We optimized UINMF and iNMF with *K* = 30, λ = 5, and took the best of 5 random initializations for 10 random seeds. We performed quantile normalization for each optimized object using the SNARE-seq dataset as a reference. Using the SNARE-seq cell labels as ground truth, we calculated the ARI and Purity scores for each of the ten seeds at louvain resolutions from 0.1 to 1.0 in increments of 0.1. We set nearest neighbors to 100 when generating the UMAPs shown, and the louvain resolution to 1. To assess the difference between UINMF and iNMF functioning, we performed a paired, one-sided Wilcoxon test between the ARI (*P =* 5.934 × 10^−15^) and Purity (*P* = 1.046 × 10^−11^) values. The UMAPs shown have a louvain resolution of 1.0, are labeled by jointly examining the original STARmap and SNARE-seq cell labels.

To assess the performance of Seurat, we used the 944 shared genes, and set the PCA dimensions equal to 30. We then calculated the Purity and ARI scores for each Louvain resolution from 0.1 to 1.0, in increments of 0.1. We conducted a paired, one-sided Wilcoxon test between UINMF and Seurat performance in terms of ARI (*P* = 0.001562) and Purity scores (*P* = 3.016 × 10−16).

### Selecting *K* and λ

The default parameter settings for iNMF are *K* = 30 and λ = 5. To allow for a fair comparison between iNMF and UINMF, we used these parameters settings for both algorithms when performing the SNARE-seq integration with intergenic peaks, the SNARE-seq integration with the 1,020 gene STARmap dataset, and the cross-species analysis. For the spatial transcriptomic dataset integrations using osmFISH (33 genes) and STARmap (28 genes, iNMF required the selected *K* to be less than the number of genes. Since one of the advantages of our algorithm is that it does not have this constraint, we selected a value of *K* = 40 for both spatial transcriptomics integrations, the largest *K* we could select without severely impacting the alignment scores (**Supplementary Fig. 2**). To select λ for the spatial transcriptomics datasets, we selected λ = (10,1) for the STARmap integration with the scRNA-seq data. The higher penalty is assigned to the STARmap data, and the use of the vectorized lambda showed improved alignment scores over 5 initializations (Supplemental Figure 3). In order to highlight that the default choice of lambda still provides statistically significantly improvement, and that the improved results were not driven by the use of a vectorized λ, we selected λ = 5 for the osmFISH dataset (**Supplementary Fig. 3**).

### Statistical Analysis

All statistical analyses were performed using R (4.0.0). P-values were calculated using one-sided paired Student T-tests or one-sided paired Wilcoxon Rank Sum Tests, as indicated. P-values are reported throughout, with statistical significance considered P < 0.05. All error bars represent standard error.

## Acknowledgments

This work was supported by NIH grants R01 AI149669, R01 HG010883, RF1 MH123199 (JDW). ARK was supported by the Genome Science Training Program T32 HG000040.

## Author Contributions

JDW conceived the idea of UINMF. JDW and ARK developed and implemented the UINMF algorithm. ARK carried out data analyses. JDW and ARK wrote the paper. All authors read and approved the final manuscript.

## Competing Interests

A patent application on LIGER has been submitted by The Broad Institute, Inc., and The General Hospital Corporation with J.D.W. listed as an inventor. The remaining authors declare no competing interests.

## Data Availability

All datasets used in this paper are previously published and freely available.

- Mouse Frontal Cortex SNARE-seq cells from Chen et al.^1^ (GSE126074)
- Mouse frontal cortex spatial transcriptomic cells from Moffit et al.^8^ (https://www.starmapresources.com/data)
- Mouse cells from the somatosensory cortex^11^ (http://linnarssonlab.org/osmFISH/)
- Adult mouse brain cells from Saunders et al.^24^(http://dropviz.org/)
- Lizard Pallium cells from Tosches, et. al^.29^ (https://public.brain.mpg.de/Laurent/ReptilePallium2018/)

## Code Availability

An R implementation of the UINMF algorithm is available as part of the LIGER package: https://github.com/welch-lab/liger

## Notes

https://github.com/welch-lab/liger

## References

1. Chen, S., Lake, B. B. & Zhang, K. High-throughput sequencing of the transcriptome and chromatin accessibility in the same cell. Nat. Biotechnol. 37, 1452–1457 (2019).

2. Liu, J., Huang, Y., Singh, R., Vert, J. P. & Noble, W. S. Jointly embedding multiple single-cell omics measurements. BioRxiv (2019).

3. Ma, S. et al. Chromatin Potential Identified by Shared Single-Cell Profiling of RNA and Chromatin. Cell 183, 1103–1116.e20 (2020).

4. Genomics, 10x. Chromium Next GEM Single Cell Multiome ATAC + Gene Expression Reagent Kits User Guide. (2020).

5. Li, G. et al. Joint profiling of DNA methylation and chromatin architecture in single cells. Nat. Methods 16, 991–993 (2019).

6. Clark, S. J. et al. scNMT-seq enables joint profiling of chromatin accessibility DNA methylation and transcription in single cells. Nat. Commun. 9, 781 (2018).

7. Method of the Year 2020: spatially resolved transcriptomics. Nat. Methods 18, 1 (2021).

8. Moffitt, J. R. et al. Molecular, spatial, and functional single-cell profiling of the hypothalamic preoptic region. Science 362, (2018).

9. Wang, X. et al. Three-dimensional intact-tissue sequencing of single-cell transcriptional states. Science 361, (2018).

10. Gyllborg, D. et al. Hybridization-based in situ sequencing (HybISS) for spatially resolved transcriptomics in human and mouse brain tissue. Nucleic Acids Res. 48, e112 (2020).

11. Codeluppi, S. et al. Spatial organization of the somatosensory cortex revealed by osmFISH. Nat. Methods 15, 932–935 (2018).

12. Richardson, S., Tseng, G. C. & Sun, W. Statistical Methods in Integrative Genomics. Annu Rev Stat Appl 3, 181–209 (2016).

13. Shen, R., Olshen, A. B. & Ladanyi, M. Integrative clustering of multiple genomic data types using a joint latent variable model with application to breast and lung cancer subtype analysis. Bioinformatics 25, 2906–2912 (2009).

14. Wang, B. et al. Similarity network fusion for aggregating data types on a genomic scale. Nat. Methods 11, 333–337 (2014).

15. Butler, A., Hoffman, P., Smibert, P., Papalexi, E. & Satija, R. Integrating single-cell transcriptomic data across different conditions, technologies, and species. Nat. Biotechnol. 36, 411–420 (2018).

16. Argelaguet, R. et al. MOFA+: a statistical framework for comprehensive integration of multi-modal single-cell data. Genome Biol. 21, 111 (2020).

17. Gayoso, A. et al. Joint probabilistic modeling of single-cell multi-omic data with totalVI. Nat. Methods (2021) doi:10.1038/s41592-020-01050-x.

18. Risso, D., Ngai, J., Speed, T. P. & Dudoit, S. Normalization of RNA-seq data using factor analysis of control genes or samples. Nat. Biotechnol. 32, 896–902 (2014).

19. Jacob, L., Gagnon-Bartsch, J. A. & Speed, T. P. Correcting gene expression data when neither the unwanted variation nor the factor of interest are observed. Biostatistics 17, 16–28 (2016).

20. Johnson, W. E., Li, C. & Rabinovic, A. Adjusting batch effects in microarray expression data using empirical Bayes methods. Biostatistics 8, 118–127 (2007).

21. Korsunsky, I. et al. Fast, sensitive and accurate integration of single-cell data with Harmony. Nat. Methods 16, 1289–1296 (2019).

22. Welch, J. D. et al. Single-Cell Multi-omic Integration Compares and Contrasts Features of Brain Cell Identity. Cell 177, 1873–1887.e17 (2019).

23. Chen, H. et al. Assessment of computational methods for the analysis of single-cell ATAC-seq data. Genome Biol. 20, 241 (2019).

24. Saunders, A. et al. Molecular Diversity and Specializations among the Cells of the Adult Mouse Brain. Cell 174, 1015–1030.e16 (2018).

25. Touzot, A., Ruiz-Reig, N., Vitalis, T. & Studer, M. Molecular control of two novel migratory paths for CGE-derived interneurons in the developing mouse brain. Development 143, 1753–1765 (2016).

26. Takata, N. & Hirase, H. Cortical layer 1 and layer 2/3 astrocytes exhibit distinct calcium dynamics in vivo. PLoS One 3, e2525 (2008).

27. Nishiyama, A., Suzuki, R. & Zhu, X. NG2 cells (polydendrocytes) in brain physiology and repair. Front. Neurosci. 8, 133 (2014).

28. Hamanaka, G., Ohtomo, R., Takase, H., Lok, J. & Arai, K. White-matter repair: Interaction between oligodendrocytes and the neurovascular unit. Brain Circ 4, 118–123 (2018).

29. Tosches, M. A. et al. Evolution of pallium, hippocampus, and cortical cell types revealed by single-cell transcriptomics in reptiles. Science 360, 881–888 (2018).

30. Belton, J.-M. et al. Hi-C: a comprehensive technique to capture the conformation of genomes. Methods 58, 268–276 (2012).

31. Kim, J., He, Y. & Park, H. Algorithms for nonnegative matrix and tensor factorizations: a unified view based on block coordinate descent framework. J. Global Optimiz. 58, 285–319 (2014).

32. Kim, J. & Park, H. Toward Faster Nonnegative Matrix Factorization: A New Algorithm and Comparisons. in 2008 Eighth IEEE International Conference on Data Mining 353–362 (2008).

