## Supplemental Figures for "Nonnegative matrix factorization integrates single-cell multi-omic datasets with partially overlapping features"

Supplementary Information

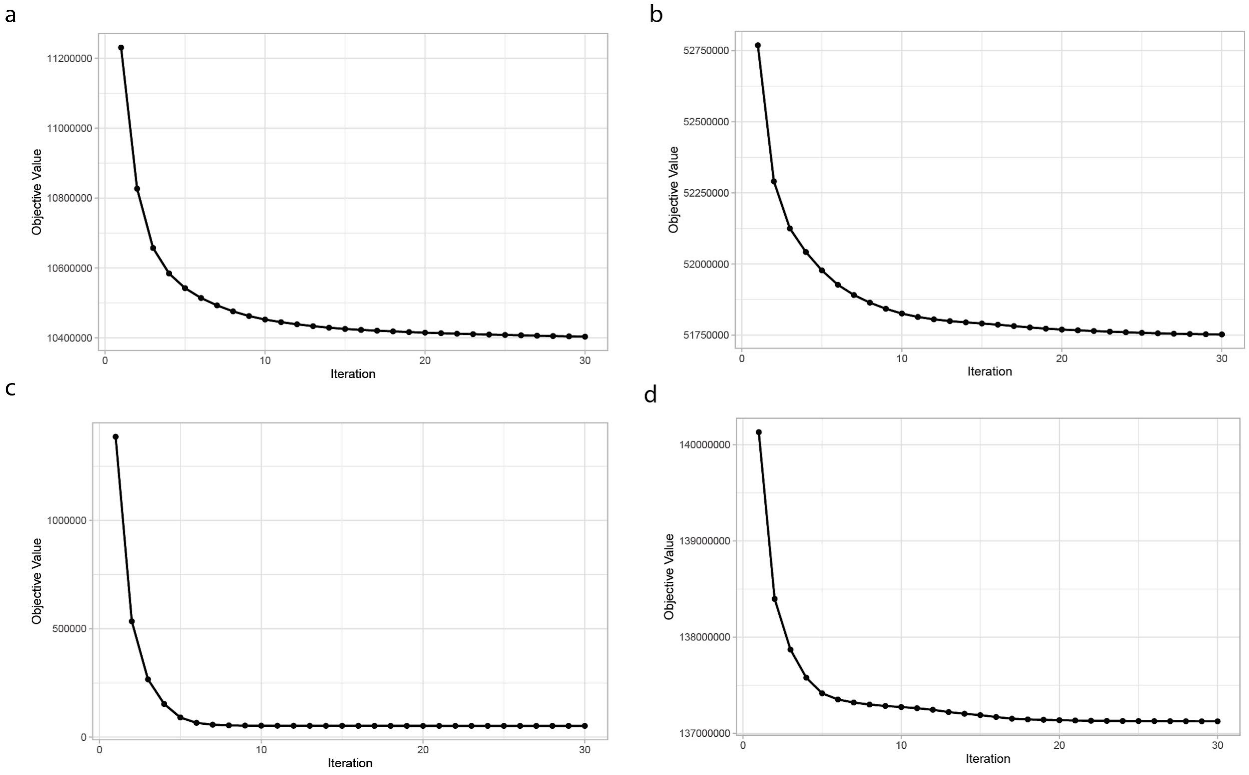


**Supplementary Figure 1. UINMF Shows Consistent Convergence of the Objective Function Values**. The objective function values for iNMF (a) and UINMF (b) across progressive iterations for the osmFISH and scRNA-seq analysis. The objective function values for iNMF (c) and UINMF (d) across progressive iterations for the SNARE-seq and STARmap data set integration. The first initialization is not shown, as it was large enough that including it resulted in the loss of the ability to visually identify the convergence of the successive objective values.

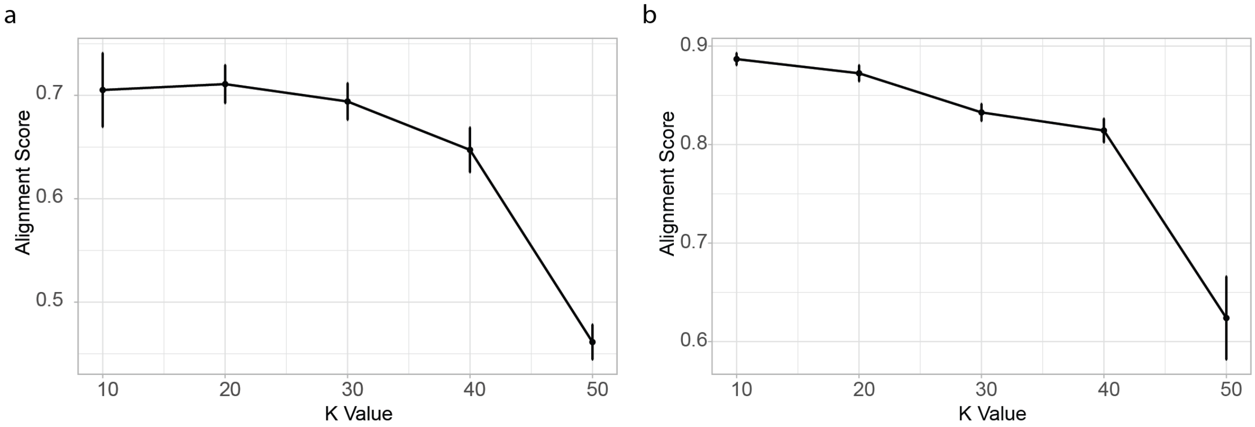


**Supplementary Figure 2. The K-value chosen has a wide range of appropriate values.** The alignment scores for the STARMAP (a) and osmFISH (b) are not severely impacted by the choice of K until an unusually large value of K (K > 40) is chosen. Alignment scores were averaged over ten initializations.

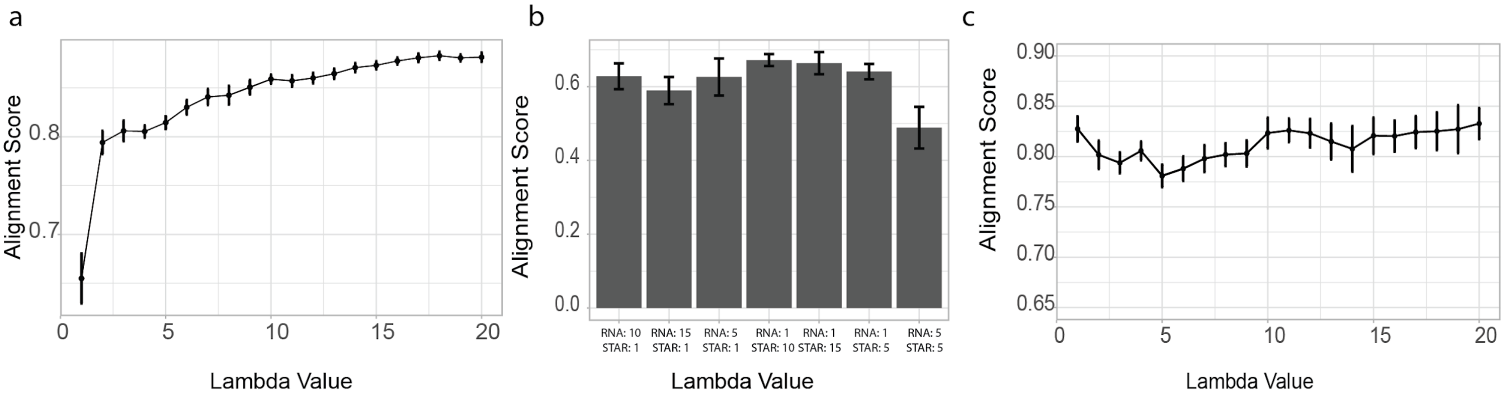


**Supplementary Figure 3. Alignment score is robust to choice of lambda.** We calculated the alignment score of the algorithm across varying values of lambda for the SNARE-seq and STARmap data integration. The results of ten initializations are shown (a). For the STARmap data, we found the penalty 10,1, where the higher penalty is assessed to the STARmap dataset, yielded the highest alignment over 5 random initializations (b). Likewise, we calculated the alignment score over 10 random initializations for a variety of lambdas for the osmFISH integration (c).

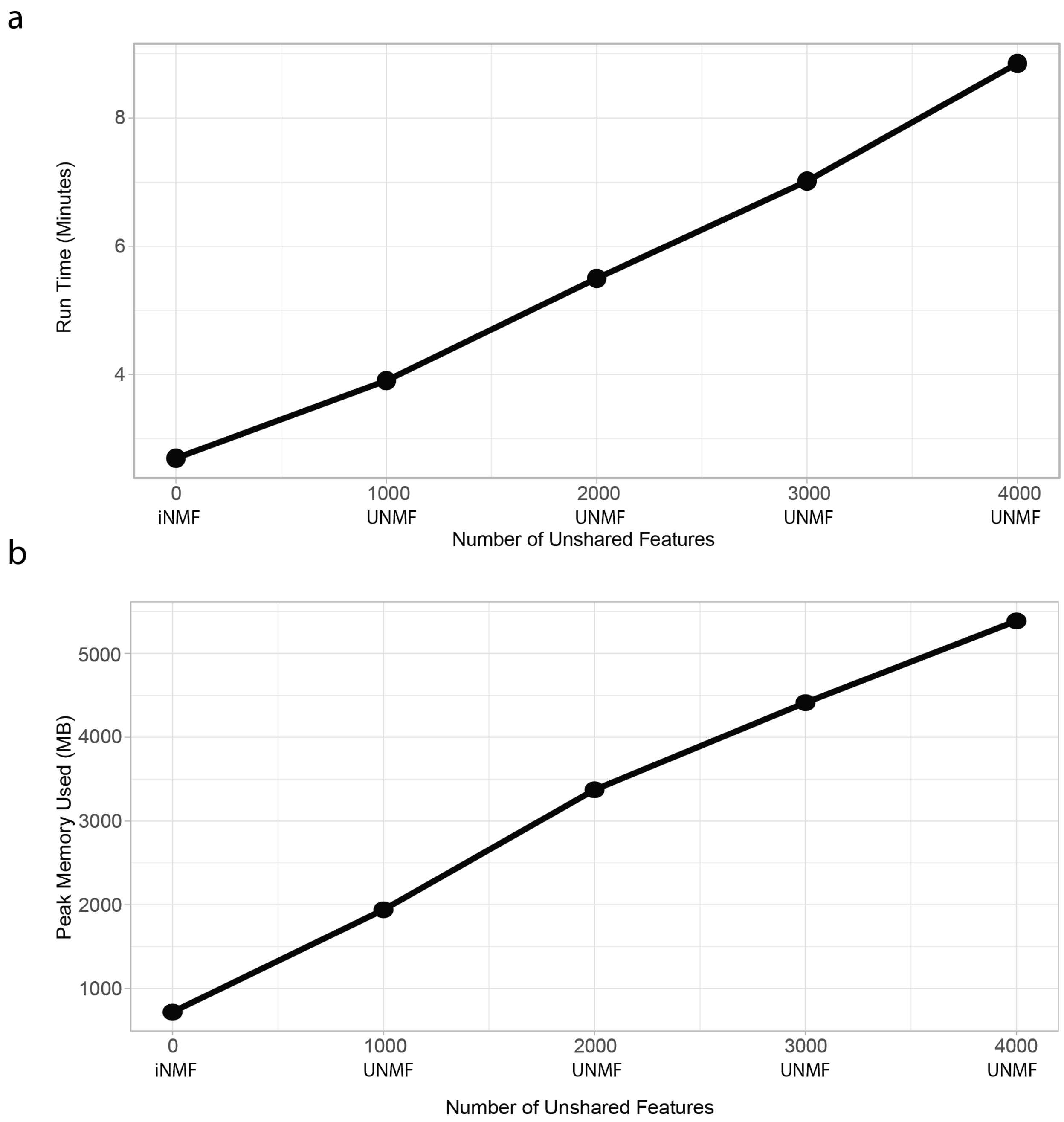


**Supplementary Figure 4. Incorporating unshared features with UINMF does not significantly increase runtime or memory usage.** Using the SNARE-seq and STARmap data, we analyze the runtime for including 0 unshared features (iNMF), as well as differing numbers of unshared features (UINMF) **(a)**. We also examine the relationship between an increasing number of features and an increase in peak memory usage **(b)**.

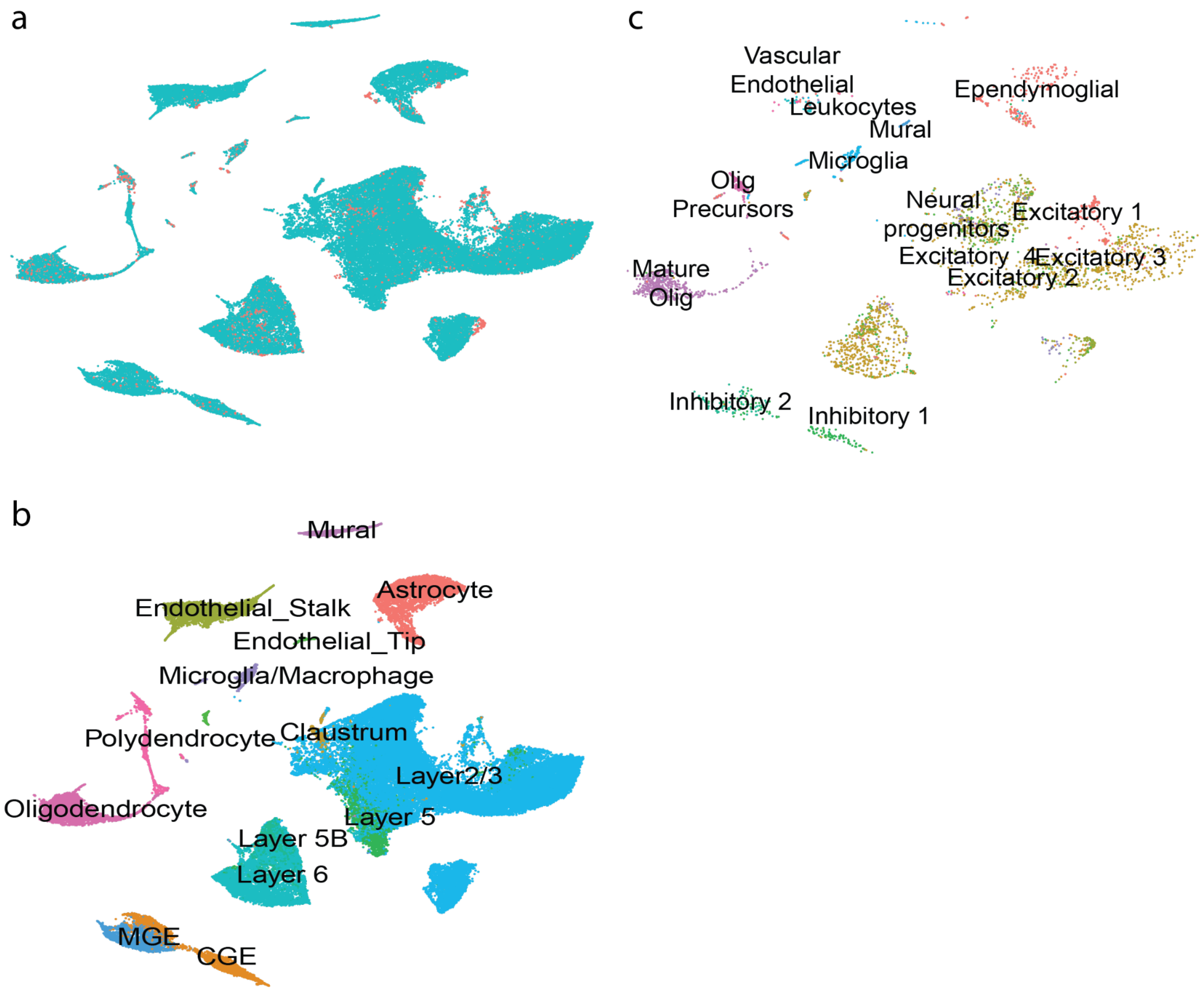


**Supplementary Figure 5. The integration of the cross-species data using only the homologous genes is not as refined.** Integrating the lizard and mouse datasets using only homologous genes, we still show adequate alignment between the two datasets **(a)**. To examine cell type correspondence between the two datasets, we examined the mouse **(b)** and lizard **(c)** cells separately, labeled with their originally published labels.
